# Single-cell spatial transcriptomics and snRNA-seq decoding the organizational principles of functional modules in the mouse amygdala

**DOI:** 10.64898/2026.06.04.730249

**Authors:** Bin Yu, Xuan Chen, Yiheng Xu, Youzhe He, Lifang Wang, Zihan Wu, Jianhua Yao, Shuqi Liu, Dan Yang, Shuxia Cao, Hao Wang, Lei Han, Xiao-Ming Li

**Affiliations:** Institute of Brain and Cognitive Science, School of Medicine, Hangzhou City University, Hangzhou, 310015, China; Affiliated Mental Health Center & Hangzhou Seventh People’s Hospital and School of Brain Science and Brain Medicine, Zhejiang University School of Medicine, Hangzhou, 310058, China; Department of Neurology and Department of Psychiatry, the Second Affiliated Hospital, Zhejiang University School of Medicine, Hangzhou, 310058, China; BGI Research, Hangzhou, 310030, China; College of Life Sciences, University of Chinese Academy of Sciences, Beijing, 100049, China; Tencent AI Lab, Shenzhen, 518057, China; NHC and CAMS Key Laboratory of Medical Neurobiology, MOE Frontier Center of Brain Science and Brain-machine Integration, and School of Brain Science and Brain Medicine, Zhejiang University School of Medicine, Hangzhou, 310058, China; Department of Neurology, Affiliated Sir Run Run Shaw Hospital, Zhejiang University School of Medicine, Hangzhou, 310058, China; Institute of Pharmacology & Toxicology, College of Pharmaceutical Sciences, Zhejiang University, Hangzhou, 310058, China

## Abstract

The amygdala is a functionally heterogeneous nuclear complex comprising multiple subnuclei that orchestrate diverse behaviors, making it essential for survival and reproduction. However, the precise neural mechanisms underlying these heterogeneous functions remain elusive, primarily due to limited knowledge of the amygdala’s cellular heterogeneity, developmental origins, spatial organization, and gene expression profiles. Here, we integrate single-cell-resolution spatial transcriptomics with precise anatomical dissection and single-nucleus RNA sequencing to systematically map the subnuclear enrichment patterns of amygdalar cell types. We further demonstrate that developmental origin determines subnuclear positioning and propose a novel functional modular architecture based on cellular composition and gene expression signatures. Furthermore, we reveal that the central amygdala (CEA) exhibits developmental heterogeneity, with *Isl1*^+^ neurons in its medial subdivision (CEAm) predominantly originating from hypothalamic progenitors. Our findings establish a spatially resolved cellular and molecular framework for investigating amygdalar functions at cellular type resolution and understanding amygdala-related neuropsychiatric disorders.

## Introduction

The amygdala represents an evolutionarily ancient and conserved brain structure, exhibiting primitive subnuclear differentiation in lungfish that becomes more organized into distinct nuclei in tetrapods, particularly mammals^1–4^. This evolutionary conservation underscores its fundamental role in survival and reproduction. Functionally, the amygdala demonstrates remarkable heterogeneity, mediating both emotional-cognitive processes including fear, anxiety and reward, as well as instinctive behaviors such as fight-or-flight responses, parenting, mating, social interaction and feeding^5–9^. Consequently, amygdala dysfunction is implicated across diverse neuropsychiatric disorders spanning neurodevelopmental conditions, psychiatric illnesses and even neurodegenerative diseases^10–13^.

The functional complexity of the amygdala, evidenced by its diverse roles in behavior and involvement in multiple neuropsychiatric disorders, stems from its heterogeneous cellular composition and extensive downstream connectivity^5, 11, 14, 15^. Recent advances in single-cell multi-omics technologies have enabled initial characterization of amygdalar cell types across species^16–21^. However, these studies remain limited by insufficient spatial resolution and have yet to systematically examine how spatial organization patterns correlate with subnuclear architecture, functional domains, developmental origins, and gene expression profiles. Elucidating these relationships is essential for further understanding the amygdala’s global connectivity patterns, functional specialization, and even evolutionary origin - all critical factors underlying its diverse physiological roles and pathological involvement in brain disorders.

This study integrates precise neuroanatomical dissection, single-nucleus RNA sequencing, and Stereo-seq spatial transcriptomics to delineate the spatial and subnuclear organization of amygdalar cell types in mice at single-cell resolution. By systematically examining the relationships between cellular architecture, developmental origins, and gene expression profiles, we establish a novel organizational framework for amygdalar subnuclei and functional modules. These findings provide a foundational basis for advancing our understanding of amygdala function and its associated neuropathologies.

## Results

### Integrating fine dissection and snRNA-seq for spatial mapping of amygdalar cell types

To investigate the spatial distribution patterns of distinct cell types across various nuclei in the mouse amygdala, we employed a strategy combining precise dissection of fresh amygdalar subregions with single-nucleus RNA sequencing (snRNA-seq). We reused the whole amygdala (Amy), basolateral complex (BLA), and central area (CEA) datasets from our previous study^16^. To better recapitulate the in vivo cellular composition and capture rare cell types, particularly given that medial nucleus (MEA) has been reported to contain exceptionally diverse neuronal subtypes^22, 23^, we additionally dissected MEA and cortical nucleus (COA) for snRNA-seq. Consequently, our mouse amygdala snRNA-seq dataset encompasses subregions including the BLA, CEA, MEA, and COA (Figures 1A and 1B; Table S1). It should be noted that the intercalated nucleus (IA) was excluded from this dissection study due to its diffuse anatomical distribution. After quality control and filtering, 73285 high-quality single nuclei from distinct dissections were integrated, revealing major cell classes via established marker expression (Figures S1A-S1C). Unsupervised clustering identified 49 transcriptionally distinct cell types, over half of which were inhibitory cell types, consistent with the known diversity of inhibitory populations in the amygdala (Figure 1C). Subsequently, cell type annotations were refined using established reference datasets and marker gene expression, enabling detection of rare cell types (e.g., Cajal-Retzius cells (CR) and chandelier cells (ChC)) through expanded nuclear sampling^24, 25^ (Figure 1C; Table S2).

**Figure 1.**
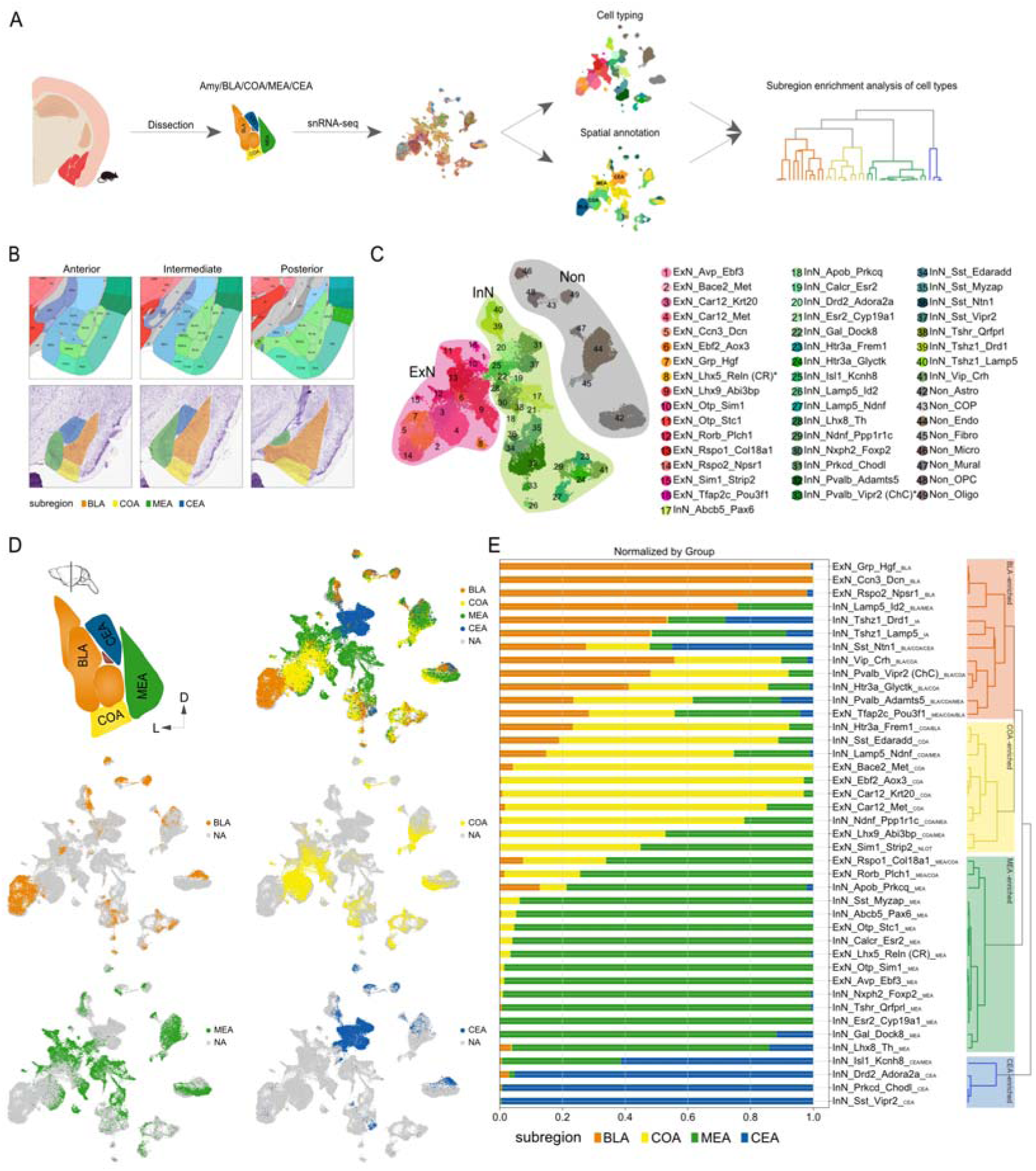
**Spatial mapping of amygdalar cell type distributions through integrated fine dissection and snRNA-seq**. (A) Workflow of mouse amygdalar subregion dissection, single-nucleus RNA-seq, and downstream analysis. (B) Anatomical schematic of mouse amygdalar subregion. Top, Allen Brain Atlas reference; Bottom, schematic of dissected subregions in corresponding coronal sections. See Table S1. (C) UMAP plot of mouse amygdalar cell types. (D) Subregion-specific enrichment of mouse amygdalar cells in UMAP space. (E) Proportional composition of mouse amygdalar cell types by anatomical subregions, and hierarchical clustering of cell types based on proportion. Major contributing subregions (>20%) labeled at bottom right of each cell type. See also Figure S1.

All nuclei retained dissection information. To determine the subregional origin of each cell type, we first visualized nuclei distribution from different subregions on UMAP. Notably, nuclei - particularly neuronal nuclei from different subregions - formed spatially segregated clusters within the UMAP embedding, reflecting both intra-subregion transcriptional homogeneity and inter-subregion transcriptional divergence (Figures 1D and S1A). These observations strongly implicate that transcriptionally distinct cell types are spatially organized within specific amygdalar subregions. To determine the subregional distribution patterns of each cell type, we quantified the proportion of nuclei from each of the four subregions within every identified cell type. As predicted, most cell types predominantly comprised nuclei enriched in specific subregions. Hierarchical clustering of proportional distribution data clearly segregated cell types into four distinct branches corresponding to BLA-, COA-, MEA-, and CEA-derived cell types (Figure 1E). These results validate, at both cellular composition and gene expression levels, the previously established amygdalar subregion classification based on anatomical features and support the functional heterogeneity across different subregions.

### Spatial distribution of ExN, InN, and their markers reveals novel subnuclear organization patterns in mouse amygdala

The dissected subregions contain multiple subnuclei. Given the technical challenges in isolating individual subnuclei, we performed Stereo-seq spatial transcriptomics^26^ on three coronal brain hemisections spanning the anteroposterior amygdala axis for each sex to systematically map amygdalar cellular spatial origins at single-cell resolution (Figure 2A; Table S1). Following data acquisition, we first delineated and excised amygdalar areas from brain slices based on anatomical landmarks (Figure 2A). Next, major cell classes were annotated using Spatial-ID, a cell typing method for spatially resolved transcriptomics via transfer learning and spatial embedding^27^. The transfer results demonstrated that neuronal cells in male and female mice exhibited significantly higher counts and detected genes than non-neuronal cells in spatial data at Bin size level (Bin 50, 50 × 50 DNB bins, 25 mm diameter) (Figure S2A). These findings are consistent with snRNA-seq data, indicating greater transcriptional activity and complexity in neurons. Our analysis of the transfer results revealed two key aspects confirming their accuracy. First, the spatial distributions of excitatory neurons (ExN), inhibitory neurons (InN), and non-neuronal cells aligned with prior knowledge: ExN predominantly localized to BLA and COA subregions, InN to CEA and MEA subregions, oligodendrocytes (oligo) to the amygdalar capsule, and endothelial/ fibroblast cells (endo/fibro) to cortical surfaces (Figure 2B). Second, ExN and InN distributions corresponded well with their respective marker gene expression patterns (Figure S2B). Notably, *Slc17a6* and *Slc17a7* showed divergent expression across amygdalar subnuclei, indicating two ExN subpopulations with distinct cellular and functional properties (Figure S2B).

**Figure 2.**
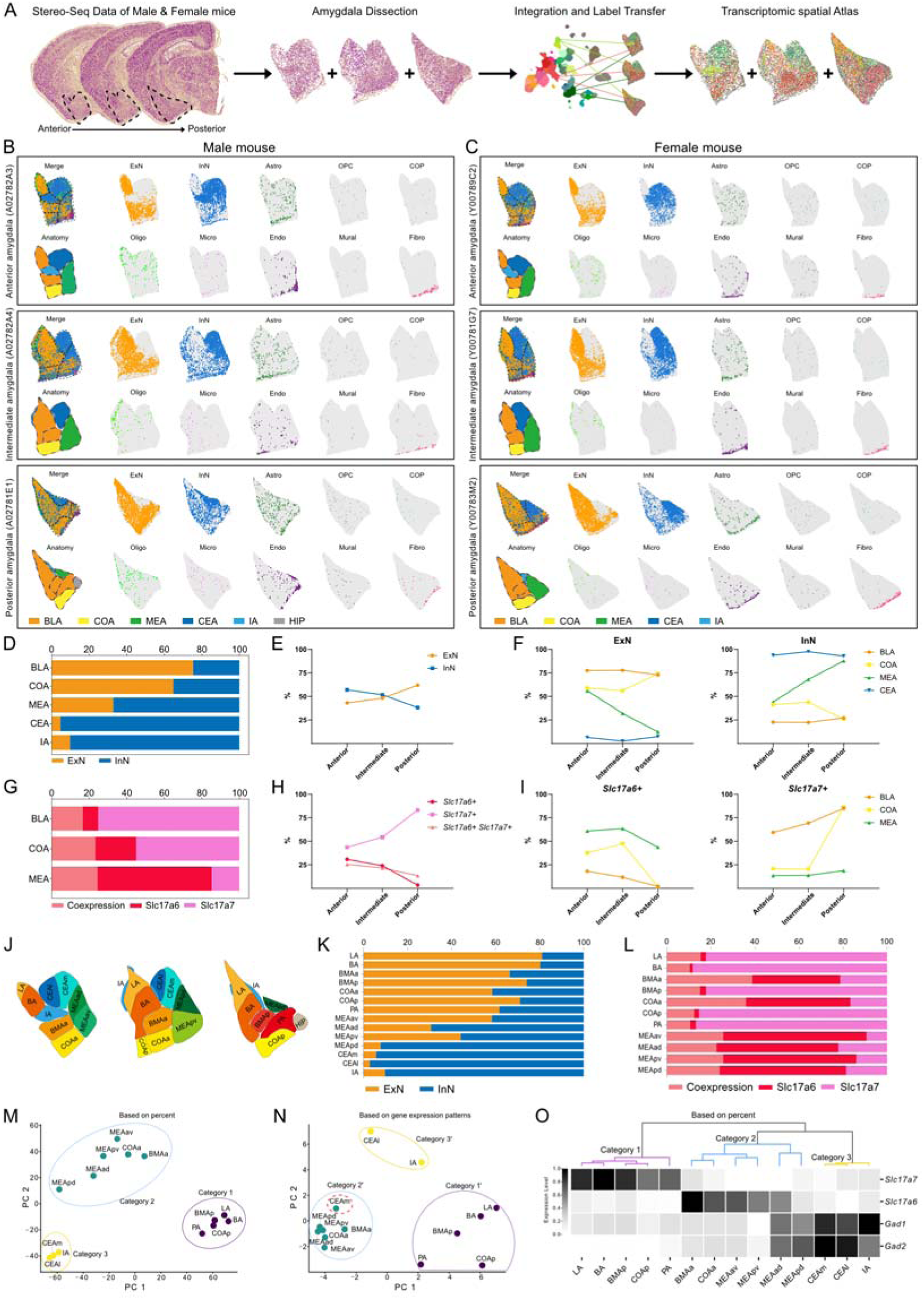
Stereo-seq spatial mapping of ExN, InN, and their markers reveals novel subnuclear organization patterns in mouse amygdala. (A) Workflow of Stereo-seq data collection and analysis. Three coronal sections spanning anterior, intermediate and posterior regions were collected for each sex. See Table S1. (B and C) Spatial distribution of cell classes in the amygdala of male (B) and female (C) mice. Dashed lines delineate subregions; chip IDs in parentheses. HIP: hippocampal region. (D) Proportions of ExN and InN across amygdala subregions (combined sexes). (E) Proportions of ExN and InN along anteroposterior amygdala axis (combined sexes). (F) Proportions of ExN and InN across subregions in anterior, intermediate, and posterior amygdala (combined sexes). (G) Proportions of *Slc17a6*^+^, *Slc17a7*^+^, and double-positive neurons across amygdala subregions (combined sexes). (H) Proportions of *Slc17a6*^+^, *Slc17a7*^+^, and double-positive neurons along anteroposterior amygdala axis (combined sexes). (I) Proportions of *Slc17a6*^+^ and *Slc17a7*^+^ neurons across subregions in anterior, intermediate, and posterior amygdala (combined sexes). (J) Schematic of the mouse amygdala subnuclear architecture. (K) Proportions of ExN and InN across amygdala subnuclei (combined sexes). (L) Proportions of *Slc17a6*^+^, *Slc17a7*^+^, and double-positive neurons across amygdala subnuclei (combined sexes). (M) PCA dimension reduction and clustering of amygdala subnuclei by ExN, InN, *Slc17a6*^+^, *Slc17a7*^+^, and double-positive neuron proportions. Colors indicate subnuclei categories. (N) PCA dimension reduction and clustering of amygdala subnuclei by feature gene expression. Colors indicate subnuclei categories’. (O) Hierarchical clustering by ExN/InN/*Slc17a6*^+^/*Slc17a7*^+^/double-positive neuron proportions, with gene expression features across subnuclei categories. Colors show normalized expression. See also Figures S2 and S3.

To characterize the spatial heterogeneity of ExN and InN across amygdalar subregions and anteroposterior axes, we performed systematic mapping of their distributions. First, spatial amygdala data were segmented into BLA, COA, MEA, CEA, and IA (main island) regions based on gene expression patterns, anatomical landmarks, and alignment with the Allen Brain Atlas^28^ (Figure 2B). We next quantified the proportions of ExN and InN populations across distinct subregions and along the anteroposterior axis. Consistent with previous reports, BLA and COA were predominantly composed of ExN, while MEA and particularly CEA/IA were mainly populated by InN (Figures 2D, S2C, and S2I). Along the anteroposterior axis, InN abundance progressively decreased from anterior to posterior regions, while ExN showed the opposite trend (Figures 2E, S2D, and S2J). Notably, subregions exhibited distinct axial gradients: COA displayed a significant posterior increase in ExN proportion, contrasting sharply with MEA where ExN decreased dramatically (posterior MEA being nearly exclusively InN). In contrast, BLA and CEA maintained relatively stable ExN/InN ratios along the axis (Figures 2F, S2E, and S2K). Given the distinct distribution patterns of *Slc17a6* and *Slc17a7* in the amygdala, we classified ExN into *Slc17a6*^+^, *Slc17a7*^+^, and *Slc17a6*^+^ *Slc17a7*^+^ populations and analyzed their subregional and anteroposterior distributions. Strikingly, We observed graded variations across BLA, COA, and MEA: BLA was predominantly *Slc17a7*^+^ ExN, COA showed decreased *Slc17a7*^+^ but increased *Slc17a6*^+^ proportions compared to BLA, while MEA exhibited an inverse pattern dominated by *Slc17a6*^+^ ExN (Figures 2G, S2F, and S2L). These ExN subtype distributions reflect functional heterogeneity among amygdalar subregions. Along the anteroposterior axis, both overall and subregion-specific analyses revealed increasing proportions of *Slc17a7*^+^ ExN and decreasing proportions of *Slc17a6*^+^ ExN. This gradient was most pronounced in COA, indicating substantial differences in cellular composition and function between its anterior and posterior portions (Figures 2H, 2I, S2G, S2H, S2M, and S2N). These patterns were also sex-independent.

Building on observed Slc17a6+/Slc17a7+ ExN axial gradients indicating intra-subregional heterogeneity, we performed finer anatomical segmentation using Allen Brain Atlas reference to resolve amygdalar organization at subnuclear resolution^28^ (Figures 2J, S3A, and S3B). We next analyzed the proportions of ExN, InN, *Slc17a6*^+^, *Slc17a7*^+^, and *Slc17a6*^+^ *Slc17a7*^+^ populations at the subnuclear level. Key findings were: 1) MEApd was predominantly InN; 2) BMAa and COAa showed relatively lower ExN proportions (∼60%); and most strikingly, 3) the *Slc17a6*^+^/*Slc17a7*^+^ profiles in BMAa/COAa closely matched MEA subnuclei but differed markedly from other BLA/COA subnuclei (Figures 2K, 2L, S3C, S3D, S3G, and S3H). To systematically characterize these patterns, we performed PCA analysis of the proportional data, revealing three distinct categories with BMAa and COAa grouping with MEA subnuclei, suggesting shared cellular and functional properties (Figures 2M, S3E, and S3I). For functional validation, we conducted gene expression-based PCA, which similarly identified three categories. While largely consistent with proportion-based clustering, CEAm showed differential grouping - clustering with CEAl/IA in proportion analysis but with MEA subnuclei in expression analysis (Figures 2N, S3F, and S3J). Together, these results demonstrate that only relative proportions of ExN, InN, and *Slc17a6*^+^/*Slc17a7*^+^ populations across amygdalar subnuclei robustly predict their functional properties, revealing three characteristic subnuclear categories. Category 1 is predominantly composed of *Slc17a7*-expressing ExN, while the category 2 shows increased representation of both *Slc17a6*-expressing ExN and InN. The third and most distinct category consists almost entirely of Gad1/Gad2-high InN (Figure 2O). This molecular stratification provides a refined framework for understanding the functional organization of amygdalar subnuclei.

### Single-cell resolved subnuclear distribution of six amygdalar cell groups revealed by Stereo-seq

We previously classified amygdalar cell types by anatomical subregions (Figure 1E), yet their fine-scale spatial distribution patterns remain uncharacterized. We first performed hierarchical clustering of ExN and InN cell types based on transcriptomic similarity. Consistent with spatial transcriptomic data (Figures 2M-2O), ExN segregated into two distinct groups: one enriched in *Slc17a7* (primarily from BLA and COA) and another expressing high *Slc17a6* levels (predominantly from MEA and COA) (Figures 3A and 3B). Notably, InN clustered into four branches, three of which were classified as MGE-, CGE-, or LGE-derived based on their distinct ganglionic eminence marker expression. The MGE/CGE-derived populations, resembling cortical interneurons (“cortical-like InN” in our previous reports^16^), expressed canonical markers (*Pvalb*, *Sst*, *Lamp5*, *Htr3a*, *Vip*) and were widely distributed across amygdalar subregions^29, 30^. In contrast, LGE-derived neurons were primarily localized to CEA/IA, representing the major inhibitory projection neurons with characteristic markers (*Prkcd*, *Sst*, *Drd2*, *Tshz1*)^31–35^. The fourth branch uniquely expressed POA (preoptic area)-related genes (e.g., Prlr)^36, 37^, primarily resided in MEA, and may share developmental and functional similarities with POA neurons (Figures 3C and 3D). Collectively, amygdalar neurons can be categorized into six distinct groups, each expressing unique signature genes (Figures S4A and S4B; Table S3).

**Figure 3.**
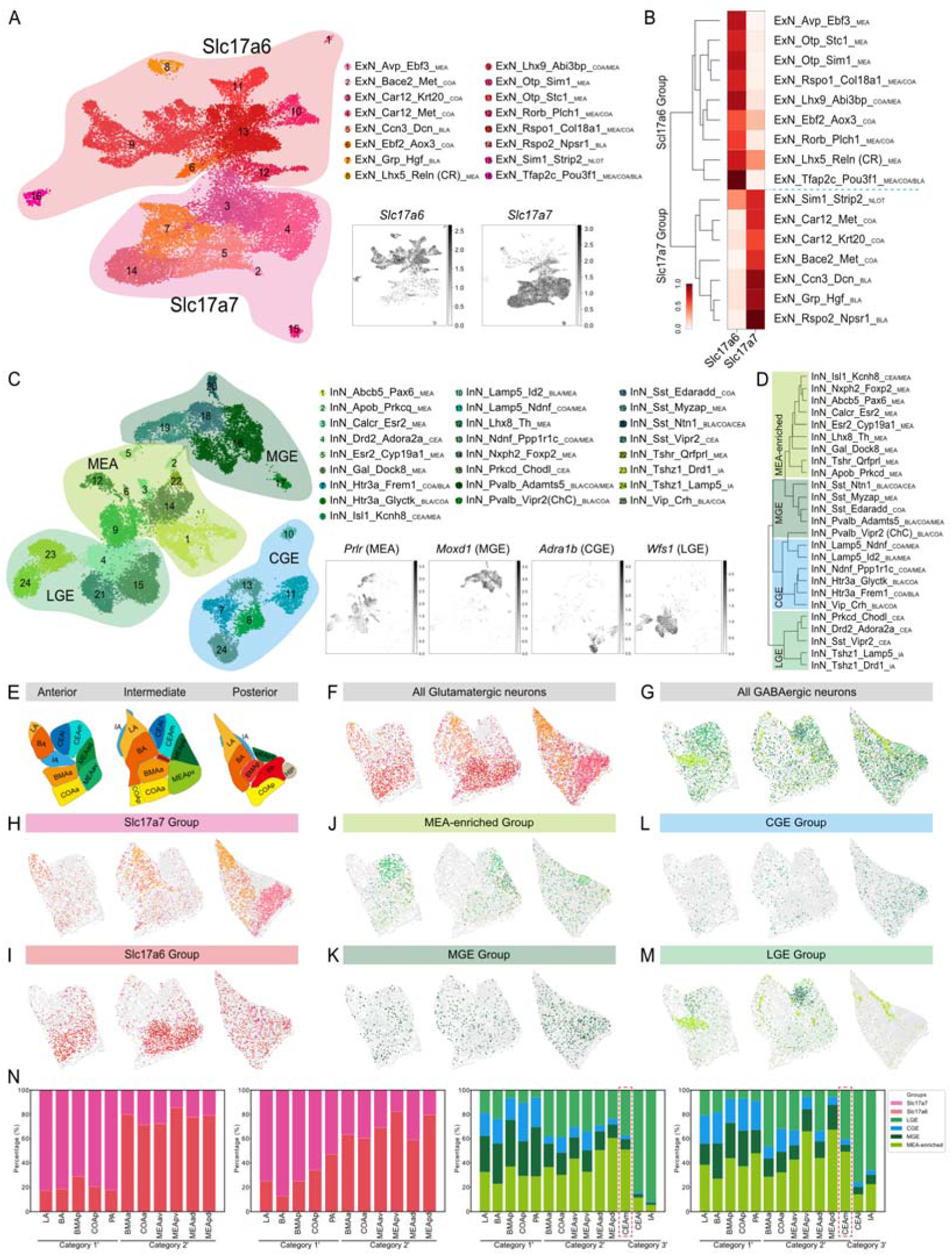
Stereo-seq unveils single-cell resolved subnuclear organization of six amygdala cell groups. (A) UMAP visualization of *Slc17a6*^+^/ *Slc17a7*^+^ excitatory neuron segregation in mouse amygdala. Insets: relative expression of *Slc17a6* and *Slc17a7*. (B) Hierarchical clustering of mouse amygdalar excitatory neurons identifies *Slc17a6*^+^ and *Slc17a7*^+^ groups, with heatmap showing normalized expression of *Slc17a6* and *Slc17a7*. (C) UMAP visualization of LGE, MGE, CGE, and MEA-enriched inhibitory neuron groups in mouse amygdala. Insets: relative expression of group marker genes. (D) Hierarchical clustering of mouse amygdalar inhibitory neurons identifies LGE, MGE, CGE, and MEA-enriched groups. (E) Schematic of the mouse amygdala subnuclear architecture. (F and G) Distribution of glutamatergic (F) and GABAergic (G) neurons across anterior, intermediate, and posterior amygdala (only male results shown). (H-M) Distribution patterns of six neuronal groups - Slc17a7 (H), Slc17a6 (I), MEA-enriched (J), MGE (K), CGE (L), and LGE (M) - across anterior, intermediate, and posterior amygdala (only male results shown). (N) Proportional distribution of six neuronal groups across amygdala subnuclei reflect PCA clustering patterns (see Figure 2N), with CEAm showing greater similarity to MEA subnuclei, BMAa and COAa than to CEAl and IA (combined sexes). See also Figures S4 and S5.

To systematically resolve the precise spatial distribution of amygdalar neuronal groups and cell types across subnuclei, we constructed a single-cell-resolution spatial atlas using the Spatial-ID transfer method (Figures S4C-F, and S5A-F; Table S4). Following Spatial-ID transfer and quality control filtering, 33598 high-quality cells were retained for subsequent analyses. Consistent with the Bin size analysis, glutamatergic neurons were predominantly distributed in BLA and COA subnuclei as well as MEAav and MEApv, while GABAergic neurons were mainly localized to IA and CEA subnuclei along with MEAad and MEApd (Figures 3E-3G). At the neuronal group level, *Slc17a7*^+^ neurons primarily occupied LA/BA/BMAp/COAp/PA (Category 1 in Figure 2M), while *Slc17a6*^+^ neurons were enriched in BMAa/COAa and MEA subnuclei (Category 2 in Figure 2M) (Figures 3H, 3I, and 3N). Among GABAergic groups, MGE- and CGE-derived cells were widely distributed across amygdalar subnuclei, with highest densities in LA/BA/BMAp/COAp/PA (Category 1’ in Figure 2N), while LGE-derived cells concentrated in IA/CEAl (Category 3’ in Figure 2N). Notably, the MEA-enriched group distributed not only in BMAa/COAa and MEA subnuclei but also abundantly in CEAm (Category 2’ in Figure 2N), mirroring transcriptome-based subnuclear clustering and suggesting CEAm may share cellular and functional properties with Category 2’ subnuclei (Figures 3J-3N). In summary, we established the most comprehensive single-cell resolution spatial atlas of the amygdala to date, delineating the relationships between inter-subnuclear gene expression similarity, cellular composition, and spatial organization of distinct cell types.

### Amygdalar neuron subnuclear distribution and functional modularity are governed by developmental origin

We have classified amygdalar cell types into six distinct groups based on gene expression profiles (Slc17a6/Slc17a7), developmental origins (MGE/CGE/LGE), and spatial distribution patterns (MEA-enriched), with detailed characterization of their subnuclear distributions. Given that amygdalar neurons originate from multiple neurogenic regions including pallium, thalamus, hypothalamus, POA, MGE, CGE and LGE^15, 31, 38–42^, we propose that these six groups reflect their distinct developmental origins. We next performed hierarchical clustering of all neurons based on transcriptomic profiles, similarly identifying six distinct groups. Notably, Groups 2, 5, and 6 specifically expressed LGE, MGE, and CGE markers respectively, consistent with their developmental origins, while Group 1 (corresponding to the previously defined Slc17a7 group) exhibited high expression of cortical marker genes (*Slc17a7*, *Neurod2*, *Emx1*)^43^, likely originating from the reported lateral pallium or dorsal pallium (LP/DP) (Figure 4A).

**Figure 4.**
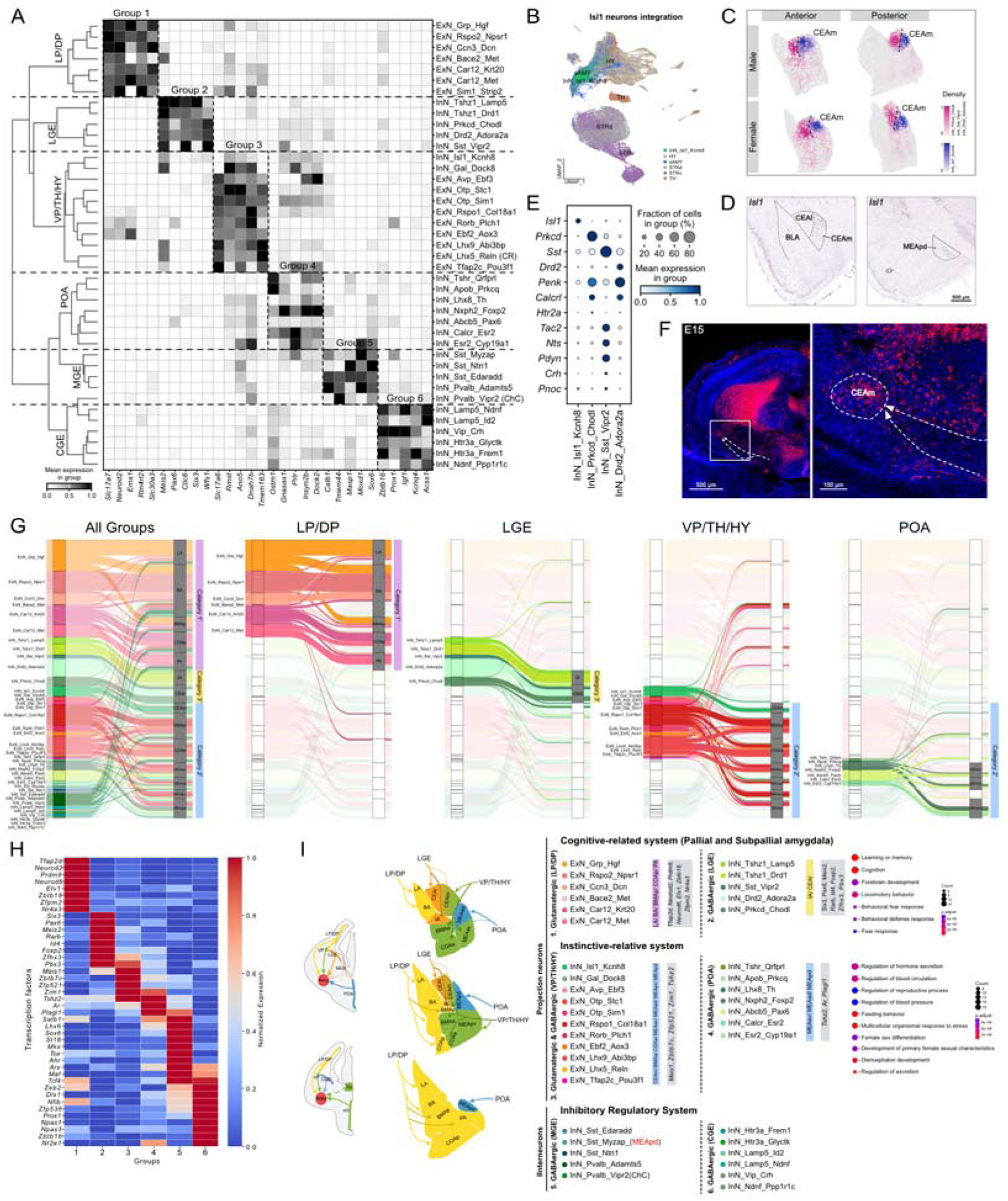
Developmental origin dictates amygdalar neuron subnuclear distribution and functional modularity. (A) Hierarchical clustering and developmental origin gene expression patterns of mouse amygdalar neuron types. (B) UMAP projection integrating InN_Isl1_Kcnh8 with published *Isl1*^+^ neurons across brain regions. (C) Spatial distribution of neuronal types in mouse central amygdala highlights InN_Isl1_Kcnh8 enrichment in CEAm. Colors indicate neuronal population densities. (D) ISH data from Allen Brain Atlas show *Isl1* expression predominantly in CEAm and MEApd of mouse amygdala.“ (E) Expression patterns of established CEA functional neuron markers across CEA neuronal types. (F) RNAscope staining shows the distribution of *Isl1*^+^ neurons in the amygdala and adjacent regions at mouse embryonic day 15 (E15), as well as the migration pathways of hypothalamic-derived *Isl1*^+^ neurons (indicated by dashed lines with arrows). (G) River plot reveals amygdalar neuron types enriched in distinct subnuclear categories by developmental origin. (H) Heatmap shows transcription factor enrichment expression across developmentally defined amygdalar neuron groups. Colors indicate normalized expression. (I) Novel functional modular architecture of mouse amygdala integrating neuronal developmental origins, subnuclear distributions, and neuronal type-specific transcription factors. See also Figure S6.

Interestingly, Groups 3 and 4 generally corresponded to the previously defined *Slc17a6*^+^ and MEA-enriched groups, but with notable differences. InN_Isl1_Kcnh8 and InN_Gal_Dock8 clustered more closely with *Slc17a6*^+^ ExN than with other MEA-enriched InN, suggesting that certain InN transcriptionally resemble *Slc17a6*^+^ ExN, potentially reflecting shared embryonic origins. Group 4 highly expressed markers of another inhibitory neurogenic zone, POA (e.g., Prlr)^36^, indicating a likely POA origin. Group 3 exhibited greater complexity, while ExN_Avp_Ebf3, ExN_Otp_Stc1, and ExN_Otp_Sim1 expressed hypothalamic-related (HY) markers (*Otp*, *Sim1*, *Avp*)^41, 42^, ExN_Lhx9_Abi3bp showed markers linked to ventral pallium (VP) (*Lhx9* and *Gdf10*)^15, 44^, and ExN_Ebf2_Plch1 specifically expressed *Pax6*, a marker often associated with thalamic (TH) identity^31, 41^ (Figure S6A). Notably, we observed that the transcriptional profiles of InN_Isl1_Kcnh8 and InN_Gal_Dock8 showed remarkable similarity to hypothalamic-derived excitatory neurons. This finding correlates with the known enrichment patterns of *Isl1* and *Gal* genes in the mouse hypothalamus (https://mouse.brain-map.org/)^45^, suggesting potential shared developmental origins or functional convergence between these distinct neuronal populations. To further validate this, we integrated these neurons with published *Isl1*^+^/*Gal*^+^ neurons from TH, HY, pallidum, striatum, and striatal amygdala^46^, revealing strong alignment with HY-derived populations (Figures 4B and S6B). Next, we performed Monocle analysis^47^ of Groups 3 and 4 with general hypothalamic (including POA) progenitor cells, identifying two developmental trajectories likely representing POA and HY lineages (Figure S6C). Thus, Group 3 appears to be a transcriptionally convergent cluster with mixed regional contributions, likely incorporating *Slc17a6*^+^ ExN from VP, TH and HY, as well as HY-derived InN. These results demonstrate that amygdalar neurons can be classified into six developmentally distinct groups by transcriptomics, explaining why subcortical ExN and InN show transcriptional similarity, including numerous VGluT2/GABA co-releasing neurons, due to shared embryonic origins. This also suggests that separate analysis of ExN and InN may obscure true transcriptomic differences, particularly in subcortical structures.

Previous studies proposed that CEA is primarily composed of LGE-derived inhibitory projection neurons^48, 49^. However, our analyses revealed that InN_Isl1_Kcnh8 - predominantly localized to the medial CEA (CEAm) and showing minimal expression of established CEA neuronal markers - originates from HY, as demonstrated by integration analysis, transcriptomic hierarchical clustering, and Allen Brain ISH^45^ also showing *Isl1* expression in CEAm and MEApd (Figures S5A, 4A-4E). RNAscope at E15 and even E18 further confirmed persistent migration of *Isl1*^+^ neurons from HY to CEAm and MEA (Figure 4F and S6D). These findings identify InN_Isl1_Kcnh8 as a novel HY-derived CEAm neuronal population with undefined function, establishing dual developmental origins for CEA: LGE-derived (predominantly CEAl) and HY-derived (primarily CEAm). This validates prior PCA clustering showing CEAm’s closer transcriptomic alignment with MEA/BMAa/COAa than CEAl/IA despite partial marker overlap (Figures 2N, S3F, and S3J), reflecting their shared developmental origin and revealing greater CEA heterogeneity than previously recognized.

We have presented a comprehensive single-cell spatial atlas of the amygdala, first elucidating inter-subnuclear gene expression relationships, then demonstrating how cell type distributions underlie these patterns, and ultimately proposing that developmentally related cell populations migrate to specific subnuclei to establish functional modules. The river plot analysis revealed distinct subnuclear distributions of neurons based on their developmental origins: LP/DP-derived neurons predominantly localized to LA, BA, BMAp, COAp, and PA (corresponding to PCA Category 1’); LGE-derived neurons primarily occupied IA and CEAl (Category 3’); VP/TH/HY-derived neurons were mainly distributed across MEA subnuclei, BMAa, COAa, and CEAm (Category 2’); while POA-derived neurons concentrated in MEA subnuclei, especially dorsal parts. Notably, MGE- and CGE-derived GABAergic interneurons showed widespread distribution throughout most amygdalar subnuclei except IA and CEA subnuclei, mirroring their cortical interneuron counterparts (Figures 2N, 4G, and S6E). Integration of our snRNA-seq data with published spatially annotated amygdalar cells confirmed the overall consistency of cell type distributions across subnuclei, despite less detailed anatomical classification in previous studies^46^ (Figures S6F, and S6G). We next identified enriched transcription factor (TF) marker genes for the six neuronal groups of distinct developmental origins, which likely regulate their embryonic development, migration, and subnuclear targeting processes (Figure 4H).

In summary, amygdalar neurons can be classified into projection neurons and interneurons. Based on developmental origins, interneurons comprise MGE- and CGE-derived GABAergic neurons, while projection neurons include: (1) LP/DP-derived glutamatergic neurons (*Slc17a7*^+^), (2) LGE-originating GABAergic neurons, (3) VP/TH/HY-derived glutamatergic/GABAergic neurons (*Slc17a6*^+^ in ExN), and (4) POA-derived GABAergic neurons - each developing under distinct transcriptional regulation that guides their migration and subnuclear targeting. Functionally, projection neurons were categorized into two systems: a cognitive-associated system (LP/DP- and LGE-derived neurons in LA/BA/BMAp/COAp/PA/IA/CEAl) and an instinctive-related system (VP/TH/HY- and POA-derived neurons in MEA subnuclei/BMAa/COAa/CEAm), with GO analysis confirming their respective behavioral and functional associations (Figure 4I).

## Discussion

In this study, we mapped the spatiotemporal origins of amygdalar cell types at single-cell resolution, systematically dissecting how gene expression profiles, developmental origins, and modular subnuclear distribution of cell types collectively shape this structurally organized yet functionally heterogeneous nuclear complex.

While previous studies have indeed investigated the spatial origins of amygdalar cell types^21^, their limited spatial resolution at the spot level could only reliably capture broad cell classes (e.g., ExN/InN), potentially introducing significant errors in fine-scale neuronal subtype identification. To overcome these limitations, we systematically integrated anatomical dissection with single-nucleus RNA-seq and single-cell resolution spatial transcriptomics, enabling precise characterization of amygdalar cell type distributions and gene expression profiles. This multimodal approach has allowed us to propose a novel functional modular organization of amygdalar subnuclei, providing a refined framework for understanding amygdalar function.

Previous studies on the function and developmental origins of the amygdala have largely been conducted within two classificatory frameworks proposed by Swanson et al., one categorizing functional and structural subregions including the basolateral complex, centromedial complex, and superficial cortical-like complex; and the other comprising systems such as frontotemporal (basolateral complex), autonomic (central extended), main olfactory (cortical-like), and accessory olfactory (medial extended)^14, 50^. Our work reveals previously uncharacterized heterogeneity in cellular composition, developmental origins, and functional specialization among subnuclei within these subregions (e.g., BMAa vs BMAp, COAa vs COAp, CEAl vs CEAm), demonstrating fundamental differences in their genetic signatures and putative functional roles. These findings necessitate a paradigm shift in amygdalar research: (1) functional investigations should adopt finer-grained subnuclear parcellation, and (2) classical subdivisions (BLA/CEA/COA) require redefinition given their intrinsic cellular and functional heterogeneity. Moving forward, a comprehensive understanding of amygdalar function will demand analysis at neuronal subtype resolution.

Current understanding holds that the CEA comprises cells originating from both dorsal and ventral LGE^31^. However, our integrated spatial transcriptomic and single-cell analyses identify a significant hypothalamic-derived cellular component within the CEA, particularly InN_Isl1_Kcnh8 neurons concentrated in the CEAm. This discovery supports a new anatomical classification: hypothalamic-derived CEA (CEAm) versus LGE-derived CEA (CEAl). The coexistence of these distinct developmental lineages provides a cellular basis for the CEA’s well-documented functional heterogeneity, encompassing both affective processes (fear learning, anxiety) and instinctive behaviors (predation, feeding)^51–53^.

## ACKNOWLEDGMENTS

We thank Li Liu and Dan Yang from the Core Facilities, Zhejiang University School of Medicine for their technical support. This work was supported by the STI2030-Major Projects (2021ZD0202700), the National Natural Science Foundation of China (NSFC) (82371525, 82090031 and 82288101), the Non-profit Central Research Institute Fund of Chinese Academy of Medical Sciences (2023-PT310-01), Nanhu Brain-computer Interface Institute (010904013), and Fundamental Research Funds for the Central Universities (2025ZFJH01-01 and 226-2024-00133).

## AUTHOR CONTRIBUTIONS

B.Y., X.C., Y.-H.X., and X.-M.L. conceptualized and designed the project. B.Y., X.C., and Y.-H.X. performed all data analyses. Y.-Z.H., L.-F.W., J.-H.Y., S.-X.C., S.-Q.L., D.Y. H.W. and L.H. helped with data collection and Stereo-seq data analyses. B.Y., X.C., Y.-H.X., and X.-M.L. wrote the manuscript. All authors edited and proofed the manuscript.

## Materials and methods

### Mouse husbandry and ethical compliance

All experimental procedures were performed in accordance with protocols approved by the Animal Advisory Committee of Zhejiang University using C57BL/6J mice. The animals were housed in the Laboratory Animal Center at Zhejiang University under controlled environmental conditions (temperature: 20±1°C; humidity: 50-60%) with a maximum of five same-sex adults per cage. Standard husbandry conditions included ad libitum access to food and water and a maintained 12-hour light/dark cycle.

For single-nucleus RNA sequencing (snRNA-seq) analysis, we collected cells from distinct amygdalar subregions using the following numbers of C57BL/6J mice: whole amygdala (n=24; 12 females/12 males), central area (CEA; n=48; 24 females/24 males), basolateral complex (BLA; n=36; 12 females/24 males), cortical nucleus (COA; n=24; 12 females/12 males), and medial nucleus (MEA; n=36; 12 females/24 males). Complete donor animal information is provided in Table S1. Whole amygdala, BLA, and CEA cells were derived from our prior work.

### Sample collection and isolation of nuclei

Eight-week-old male and female C57BL/6J mice were deeply anesthetized via intraperitoneal injection of 1% sodium pentobarbital (0.1 g/kg body weight) and transcardially perfused with ice-cold, carbogenated (95% O_2_/5% CO_2_) NMDG-based artificial cerebrospinal fluid (NMDG-ACSF) containing the following composition (in mM): 93 NMDG, 2.5 KCl, 1.25 NaH_2_PO_4_, 30 NaHCO_3_, 20 HEPES, 25 glucose, 5 sodium ascorbate, 2 thiourea, 3 sodium pyruvate, 10 MgSO_4_·7H_2_O, and 0.5 CaCl_2_·2H_2_O. Following decapitation, whole brains were immediately immersed in carbogenated NMDG-ACSF maintained at 0-4°C. Coronal sections (300 μm thickness) were prepared using a vibratome (VT1200s, Leica) in chilled NMDG-ACSF with continuous carbogenation. Under microscopic guidance (SMZ745, Nikon), COA and MEA were microdissected from sequential sections, flash-frozen in liquid nitrogen, and stored at-80°C until processing. All dissections were performed during the light phase of the 12:12 h light-dark cycle to control for circadian variability. Whole amygdala, BLA, and CEA samples were derived from our prior work.

Nuclei isolation was performed following the 10× Genomics protocol (CG000393 • Rev A) with modifications. Briefly, frozen tissues were homogenized on ice in lysis buffer containing the following components: 10 mM Tris-HCl (pH 7.4), 10 mM NaCl, 3 mM MgCl₂, 0.01% NP-40 (Thermo Fisher Scientific), 1 mM β-mercaptoethanol, and 0.2 U/μL RNase inhibitor (Promega) using a glass homogenizer. Homogenization was performed until complete tissue dissociation (no visible clumps). The homogenate was incubated on ice for 5 min, followed by addition of an equal volume of HEB medium (Hibernate E/B27/GlutaMAX, Gibco) to terminate lysis. The lysate was filtered through a 30-μm cell strainer (pluriSelect) to remove debris and large aggregates, then centrifuged at 500 × g for 5 min at 4°C. For additional debris removal, the Debris Removal Solution (Miltenyi Biotec) was applied according to the manufacturer’s protocol. Isolated nuclei were washed twice in wash/resuspension buffer (1× PBS, 1% bovine serum albumin (BSA, Sigma-Aldrich), and 0.2 U/μL RNase inhibitor), with centrifugation at 500 × g (4°C, 5 min) between washes. Nuclear integrity and concentration were assessed by DAPI staining (Thermo Fisher Scientific) and manual counting using a fluorescence microscope (BX53, Olympus). Finally, nuclei were diluted to a working concentration of 500–1,000 nuclei/μL in preparation for 10× Chromium single-nucleus capture.

### SnRNA-seq library preparation and sequencing

Single-nucleus RNA sequencing libraries were prepared using the Chromium Single Cell 3L GEM, Library & Gel Bead Kit v3 (10x Genomics, #1000075) following the manufacturer’s protocol (CG000204 Rev D). The workflow included: (1) nuclear suspension loading onto Chromium Next GEM Chips, (2) nuclear barcoding via gel bead-in-emulsion formation, (3) cDNA amplification, and (4) library construction. All libraries were assessed for quality by Agilent 4200 TapeStation analysis prior to sequencing. Final libraries were paired-end sequenced (150 bp × 2) on an Illumina NovaSeq 6000 platform (Illumina) at Abiosciences (Analytical Biosciences Limited, Beijing, China).

### SnRNA-seq data processing

The raw snRNA-seq data were initially processed with the Cell Ranger v7.1.0 (10x Genomics) pipeline using default parameters, and aligned to the mouse mm10 reference genome. Subsequently, the raw count data were further processed with the Scanpy (v1.9.1) package^54^. Nuclei expressing fewer than 200 genes or genes present in fewer than three nuclei were excluded from the analysis. Potential doublets were identified and filtered out using Scrublet (v0.2.3)^55^, with a threshold set at ‘scrublet_score < 0.3’. Additionally, nuclei with more than 10,000 detected genes or total counts per cell greater than 50,000 were excluded. Nuclei in which mitochondrial gene counts exceeded 2% were also removed. Following clustering, any clusters that displayed high expression of markers from multiple cell types were considered doublets and removed from further analysis. After applying stringent quality control analysis, 73,285 nuclei remained in the dataset, with a median of 4,286 unique molecular identifiers (UMIs) and 2,108 detected genes. The raw count matrix was normalized and log-transformed using the normalize_total and log1p functions in Scanpy, with default parameters. Highly variable genes (HVGs) were identified through the scanpy.pp.highly_variable_genes function with parameters: “min_mean = 0.0125, max_mean = 3, min_disp = 0.5”, and all HVGs were selected for downstream analysis.

### Dimensionality reduction, batch correction and unsupervised clustering

We utilized the Scanpy pipeline for dimensionality reduction and neighborhood graph embedding. We first performed principal component analysis (PCA) using the scanpy.tl.pca function, with parameters svd_solver =‘arpack’, n_comps = 100, which resulted in a matrix of 100 principal components. To correct batch effects between different samples, we applied the bbknn function with batch_key =’sample’, followed by the construction of the neighborhood graph. For embedding the graph in two dimensions, Uniform Manifold Approximation and Projection (UMAP) was implemented using the pp.neighbors function (parameters: n_neighbors = 15, knn = True, use_rep = ‘X_pca’ and method = ‘umap’) followed by the tl.umap function (with default parameters). Unsupervised clustering was realized using the sc.tl.leiden function with appropriate resolution.

### Marker gene selection and cell type annotation

At the Major Class level, glutamatergic and GABAergic neurons were annotated based on the expression of well-established canonical markers, such as *Slc17a6*/*Slc17a7* for glutamatergic neurons and *Gad1*/*Gad2* for GABAergic neurons, respectively. All other remaining clusters were classified as non-neuronal cells. At the Celltype level, COSG (v1.0.1)^56^ was first used to identify marker genes for each cell type with the parameters ‘mu=100, expressed_pct=0.1’. Subsequently, 49 distinct cell types were annotated through reference mapping to published mouse amygdala snRNA-seq datasets^16^, in conjunction with the expression of identified cell type-specific markers.

### Stereo-seq

#### Stereo-seq sample preparation and sequencing

For Stereo-seq, 10-µm coronal frozen section of mouse hemi-brain were cut and attached to the Stereo-seq capture chip (1 cm x 1 cm), followed by incubation at 37 °C for 3 min. The sections were subsequently fixed with methanol at −20 °C for 40 min, and then stained with a nucleic acid dye (Thermo Fisher, Q10212) to determine the location of the nuclei. The tissue slices on the chip were transduced, and the released RNA was captured by DNA nanospheres (DNBs) on the chip, followed by in situ reverse transcription to produce cDNA. For library construction, 20 ng of cDNA was used for fragmentation and amplification. Finally, the purified polymerase chain reaction (PCR) products were used for DNB production, and the libraries were sequenced on a MGI DNBSEQ-Tx sequencer.

#### Stereo-seq data processing

Raw Stereo-seq data processing was performed using previously described methods. The Stereo-seq Analysis Workflow (SAW) v6.1.3 (https://github.com/STOmics/SAW) was employed to analyze the raw data. The coordinate identity (CID) and molecular identifier (MID) were extracted from Read 1, while cDNA sequences were included in Read 2. Pre-processed reads were aligned to the mouse genome reference using STAR v2.7.6.^57^ Finally, exonic reads with CID information were used to generate an expression matrix, with gene features represented by rows and DNBs represented by columns.

#### Cell segmentation

Cell segmentation for Stereo-seq data was performed as previously described. The total UMI counts within each DNB were summed to create a spatial density matrix, which was transformed into an image where each pixel represented a DNB and its intensity reflected the UMI count. Segmentation was carried out using a pre-trained ESPANet model (https://arxiv.org/abs/2105.14447) based on the nucleic acid-stained images. A watershed algorithm was used for post-processing to generate binary masks for each cell. UMIs from DNBs within each mask were aggregated by gene, creating a cell-by-gene matrix (cellbin data) for further analysis.

### Quality control of single-cell Stereo-seq data

We implemented stringent quality control measures to ensure data reliability. Cells were excluded based on the following criteria: (1) those expressing fewer than 100 detected genes across all cell types, and (2) those exhibiting mitochondrial gene content exceeding 5% of total transcripts. These thresholds were established through systematic evaluation of gene detection distributions and mitochondrial RNA proportions across the dataset. Following this quality filtration pipeline, our final high-quality dataset contained 33598 single cells expressing average 465 genes, which were used for all downstream analyses.

### Spatial-ID transfer

To annotate cells in Stereo-seq data, we used the Spatial-ID v1.1.1 algorithm to integrate snRNA-seq and Stereo-seq data for cell-type transfer within a spatial context. A four-layer deep neural network (DNN) was first trained on common genes between the two datasets to predict initial cell type probabilities for each Stereo-seq cell. Spatial neighborhood information was encoded in an adjacency matrix, and a graph convolution network (GCN) was applied, consisting of a deep autoencoder, variational graph autoencoders, and a classifier. The autoencoders learned spatial and gene expression features, while the classifier aligned these features to cell types. The model was trained for 200 epochs with self-supervised and supervised losses, refining cell type probabilities based on both spatial and gene expression data. Final cell type annotations were determined by the highest GCN-generated probability for each cell. Spatial-ID effectively aligned cell types and reduced batch effects across Stereo-seq sections.

### Dendrogram analysis

Hierarchical clustering was performed using the sc.tl.dendrogram function in Scanpy. First, the hierarchical tree was constructed by grouping the data based on the ‘Celltype’ annotation, using the PCA-reduced representation stored in X_pca. The sc.tl.dendrogram function was applied with Pearson correlation as the distance metric (cor_method=‘pearson’), and the linkage method was set to Ward’s method (linkage_method=‘ward’) to minimize within-cluster variance.

### Idetification of *Slc17a6*^+^/*Slc17a7*^+^ populations

To classify excitatory neurons in our Stereo-seq data (bin 50) according to their vesicular glutamate transporter (*Slc17a6* and *Slc17a7*) expression profiles, we developed a quantitative analytical approach. Initial ExN populations were identified from the

Stereo-seq dataset using Spatial-ID, with raw UMI counts for *Slc17a6* and *Slc17a7* extracted from the count matrix. For each spot expressing at least one of these markers, we calculated the relative contribution ratio:

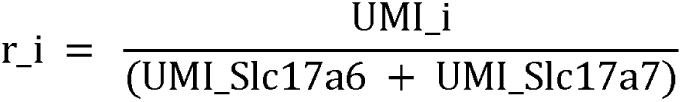

where i represents either *Slc17a6* or *Slc17a7*. To mitigate potential artifacts from spatial transcriptomic technical limitations including gene diffusion and detection sensitivity, we established stringent classification thresholds: spots with r_*Slc17a6* ≥ 0.70 were designated as *Slc17a6*^+^ population, those with r_*Slc17a7* ≥ 0.70 as *Slc17a7*^+^ population, spots showing both ratios < 0.70 as coexpression, and spots lacking detectable expression of both markers as neither population.

### Spatial registration and amygdala subnuclear segmentation

Stereo-seq data (bin 50) were registered to the Allen Mouse Brain Common Coordinate Framework version 3 (CCFv3; RRID: SCR_002978, http://atlas.brain-map.org/)through a three-step process: (1) selection of coronal reference plates from the CCFv3 atlas matching our Stereo-seq sections (A02782A3 corresponds to slice 652, A02782A4 corresponds to slice 692, A02781E1 corresponds to slice 755, Y00789C2 corresponds to slice 640, Y00781G7 corresponds to slice 688, Y00783M2 corresponds to slice 742); (2) 2D rigid registration using Leiden-clustered spots (bin 50) as spatial landmarks, with alignment optimized based on major anatomical boundaries; and (3) manual refinement of local anatomic discrepancies using Affinity Designer 2 software. Following CCFv3 registration, we performed amygdala-specific structural refinement by first removing extraneous non-amygdalar structures to focus the analysis. Related subnuclei were then consolidated by merging CEAl with CEAc (collectively designated as CEAl), grouping BLAa, BLAp and BLAv as BA, and combining COApl and COApm as COAp. Through this process, the amygdala was systematically divided into 14 distinct subnuclear compartments: LA, BA, COAa, COAp, BMAa, BMAp, PA, CEAl, CEAm, IA, MEAad, MEAav, MEApd, and MEApv.

### PCA analysis

We developed an analytical pipeline to investigate the organizational principles of amygdalar subnuclei based on ExN and InN composition. The analysis incorporated Stereo-seq data (bin 50) from three male and three female slices, processed as a combined Seurat object^58^. Cells were assigned to 14 anatomically defined subnuclei (LA, BA, BMAa, BMAp, PA, COAa, COAp, MEAav, MEAad, MEApv, MEApd, CEAm, CEAl, IA) for regional characterization. For each subnucleus, we constructed five-dimensional feature vectors comprising the ExN/InN ratio, and the relative proportions of three molecularly defined ExN subpopulations (*Slc17a6*^+^, *Slc17a7*^+^, and coexpressing cells). Principal component analysis (PCA) was performed on the resulting subnuclei × feature matrix using the prcomp function, with the first two principal components subsequently employed for k-means clustering (k=3; random seed=123). Cluster assignments were visualized in two-dimensional PCA space. To investigate potential sex differences, we repeated this analytical pipeline separately for male and female datasets by creating sex-specific Seurat objects while maintaining all other parameters constant.

To investigate subnuclear organization based on transcriptional profiles, we analyzed the same six-sample Stereo-seq dataset after assigning cells to the 14 predefined subnuclei. Using the Seurat function FindAllMarkers with stringent thresholds (min.pct = 0.3, logfc.threshold = 0.5), we identified the top five most upregulated genes (by log2 fold change) as signature genes for each subnucleus. Subnuclear-specific expression profiles were generated using AverageExpression (return.seurat = TRUE), creating a gene × subnuclei matrix that was Z-score normalized prior to principal component analysis (prcomp). We then performed k-means clustering (k = 3, random seed = 123) on the first two principal components, with results visualized as PC score plots colored by cluster assignment. To assess potential sex differences, we repeated this analytical pipeline identically for male and female datasets by creating sex-specific Seurat objects while maintaining all parameters and thresholds constant.

### RNAscope

For RNAscope multiplex fluorescence in situ hybridization (mFISH) analysis, E18 mouse brains were processed as follows: Mice were perfused with ice-cold 1× PBS, after which brains were dissected, embedded in optimal cutting temperature compound (OCT; Sakura), flash-frozen on dry ice, and stored at-80°C. Coronal sections (15-20 μm thickness) were cut using a Cryostat (CM1950; Leica) and mounted on Superfrost Plus slides (Epredia). Sections were air-dried at-20°C and stored at-80°C until processing. RNAscope mFISH was performed according to the manufacturer’s protocol (Advanced Cell Diagnostics) using the *Isl1*-specific probe (Mm-Isl1; Cat# 451931-C1). Imaging was conducted using an Olympus FV3000 confocal microscope with 10× and 20× air objectives, and subsequent image analysis was performed using ImageJ software (NIH).

### Intergation with reported snRNA-seq data

To investigate the developmental origins of specific amygdalar neuronal types, we implemented a targeted integration approach combining our single-cell RNA sequencing data with specific subsets of the Allen Brain Cell Atlas WMB-10x mouse whole-brain dataset^46^. For the InN_Isl1_Kcnh8 analysis, we selected *Isl1*-positive cells from hypothalamic (HY), thalamic (TH), striatal dorsal/ventral (STRd/STRv), and striatal amygdala (sAMY) regions in the public dataset for integration with our InN_Isl1_Kcnh8 cells. Similarly, for the InN_Gal_Dock8, we integrated *Gal*-positive cells from HY, TH, sAMY, and pallidum (PAL) regions with our corresponding neuronal population. To address sample size imbalance (2,071 InN_Gal_Dock8 cells vs. abundant public dataset *Gal*^+^ cells), we randomly subsampled 20% of the public *Gal*^+^ cells prior to integration. All data underwent consistent preprocessing: log normalization (scale factor = 10,000), selection and scaling of 3,000 highly variable genes, followed by PCA dimensionality reduction. We performed batch correction using Harmony (v1.2.0)^59^ with brain region as the covariate (InN_Isl1_Kcnh8: θ = 0.3, dims 1-9; InN_Gal_Dock8: θ = 0.9, dims 1-8). The resulting integrated low-dimensional embeddings were used for all downstream analyses.

To comprehensively characterize amygdalar cell types and their spatial distrbution, we integrated our single-cell transcriptomic data with published datasets^46^ containing amygdalar cell populations from distinct subnuclear or subregional structures. The integration was performed using Seurat (v4.3.0) through the following steps: First, we identified integration anchors using the “FindIntegrationAnchors” function, which enabled robust alignment of datasets while preserving biological variation. The integrated data were then projected into a unified UMAP space using the first 15 principal components (dims = 1:15) as input. Subsequent clustering analysis employed the “FindNeighbors” and “FindClusters” functions (resolution = 1.3) to generate an initial cellular taxonomy. This preliminary clustering was further refined through manual curation based on two key criteria: (1) systematic evaluation of canonical marker gene expression patterns across clusters, and (2) careful consideration of spatial distribution patterns within amygdalar subnuclei.

We developed a co-occurrence analysis pipeline to quantitatively evaluate the transcriptional similarity between neuronal types identified in our study and those annotated in the Allen Brain Cell Atlas. Following integration and clustering, we systematically analyzed cell type correspondence by calculating pairwise co-occurrence ratios. For each neuronal type pair between datasets, we first determined the proportion of cells assigned to each integrated cluster, then computed the minimum proportion for each cluster as its co-occurrence score, and finally summed these minimum values across all clusters to derive the final co-occurrence ratio (ranging theoretically from 0, indicating no overlap, to 1, representing complete overlap). The proportion matrix was normalized to a 0-1 scale and visualized as a grayscale heatmap using the seaborn.heatmap function (cmap=“Greys”).

### River plot

We integrated Stereo-seq datasets from six mouse amygdala slices (three male and three female slices) into a unified Seurat object. For subsequent analysis, we quantified: (1) the distribution proportions of each neuronal type across subnuclei, and (2) the cellular composition of each subnuclei by neuronal types. Proportions below 5% were excluded from analysis, retaining only those with frequency ≥5% for river plot visualization. All computational analyses were performed in R (v4.2.1) utilizing the dplyr package (v1.1.4), with river plots generated using ggalluvial (v0.12.4).

### Monocle analysis

To trace the developmental origins of VP/TH/HY (ExN_AVP_Eb3, ExN_Ebf2_AOx3, ExN_Lhx5_Reln, ExN_Lhx9_AbI3DP, ExN_Otp_Sim1, ExN_Otp_Ste1, ExN_Rorb_Pich1, ExN_Rspo1_Col18a1, ExN_Tap2c_Pou3f1, InN_Gal_Dock8, InN_Isl1_Kcnh8) and POA (InN_Abcb5_Pax6, InN_Apob_Prkcg, InN_Calcr_Esr2, InN_Esr2_Cyp19a1, InN_Lhx8_Th, InN_Nxph2_Foxp2, InN_Tshr_Qrfprl) neuronal groups, we performed pseudotemporal ordering analysis by integrating these cells with hypothalamic progenitor cells from GSE132730^60^. Both datasets were log-normalized (scale factor=10,000), and 3,000 highly variable genes were selected, centered, and scaled prior to PCA. The first eight principal components were input into “Harmony” (θ = 0.9, dims = 1 – 8) for batch correction to obtain a low dimensional embedding. The integrated data were analyzed in Monocle3 (v1.3.4)^47^ using learn_graph (minimal_branch_len=8, geodesic_distance_ratio=0.5) for trajectory inference, with root nodes manually specified in order_cells to reconstruct developmental paths from progenitors to mature neuronal states.

### Identification of transcription factors

Neuronal group-specific marker genes were identified using COSG (groupby=’group’, n_genes_user=100, mu=300, expressed_pct=0.1, remove_lowly_expressed=True). From the top 100 marker genes, we extracted transcription factors for further analysis. The AnnData object was converted to a pandas DataFrame, and expression values were averaged within each group to create a summary matrix. Visualization was performed using Seaborn (vmin=-1, vmax=1) with a coolwarm color palette, generating a heatmap that displays the average expression patterns of selected transcription factors across the defined cell-type groups.

## Data availability

The raw sequencing data have been deposited in the Genome Sequence Archive^61^ in the National Genomics Data Center^62^, China National Center for Bioinformation / Beijing Institute of Genomics, Chinese Academy of Sciences (GSA: CRA027757) and are publicly available as of the date of publication. Analyzed data are accessible through download link in GitHub repository (https://github.com/coffeei1i/amg). Additional information required to reproduce the analyses or perform reanalyses is available from the lead contact upon reasonable request.

## Code availability

The complete bioinformatics analysis pipeline and all custom scripts developed for this study have been deposited in a publicly accessible GitHub repository (https://github.com/coffeei1i/amg). All data were analyzed with standard programs and packages, as detailed in the KEY RESOURCES TABLE.

---

**Figure S1.**
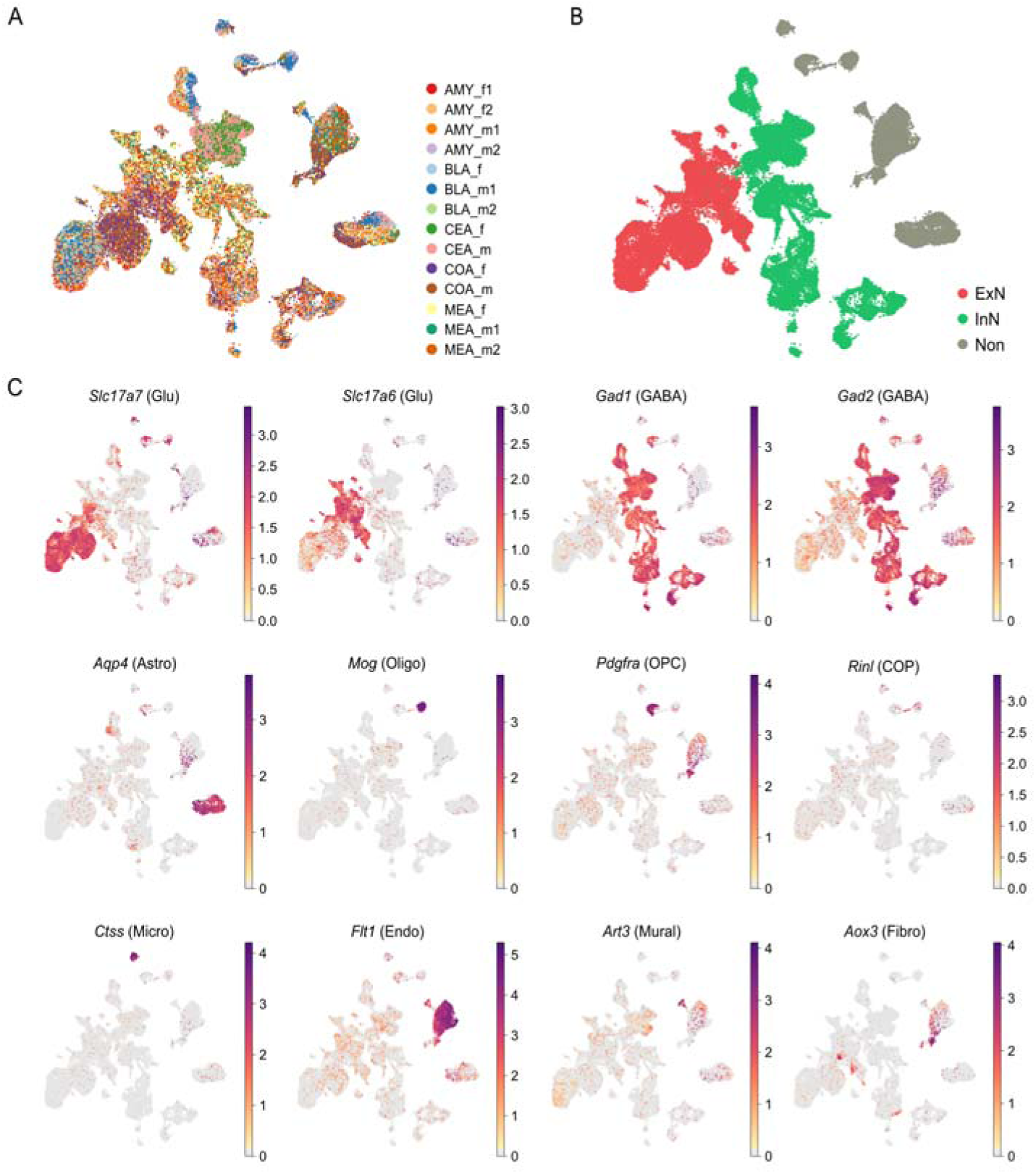
Mouse amygdala snRNA-seq data integration and visualization of marker gene expression, related to. Figure 1. (A) UMAP plot showing the integration of mouse amygdala cells from different genders and different anatomical subregions. f, female; m, male; numbers indicate sample IDs. (B) Distribution of ExN, InN, and non-neuronal cells (Non) in UMAP space. (C) UMAP visualization of marker gene expression in the snRNA-seq dataset of the mouse amygdala. Astro: astrocytes; COP: committed oligodendrocyte precursors; Endo: endothelial cells; Fibro: Fibroblasts; GABA: GABAergic neurons; Glu: Glutaminergic neurons; Micro: microglia; Mural: mural cells; Oligo: oligodendrocytes; OPC: oligodendrocyte precursor cells. Colors represent relative expression.

**Figure S2.**
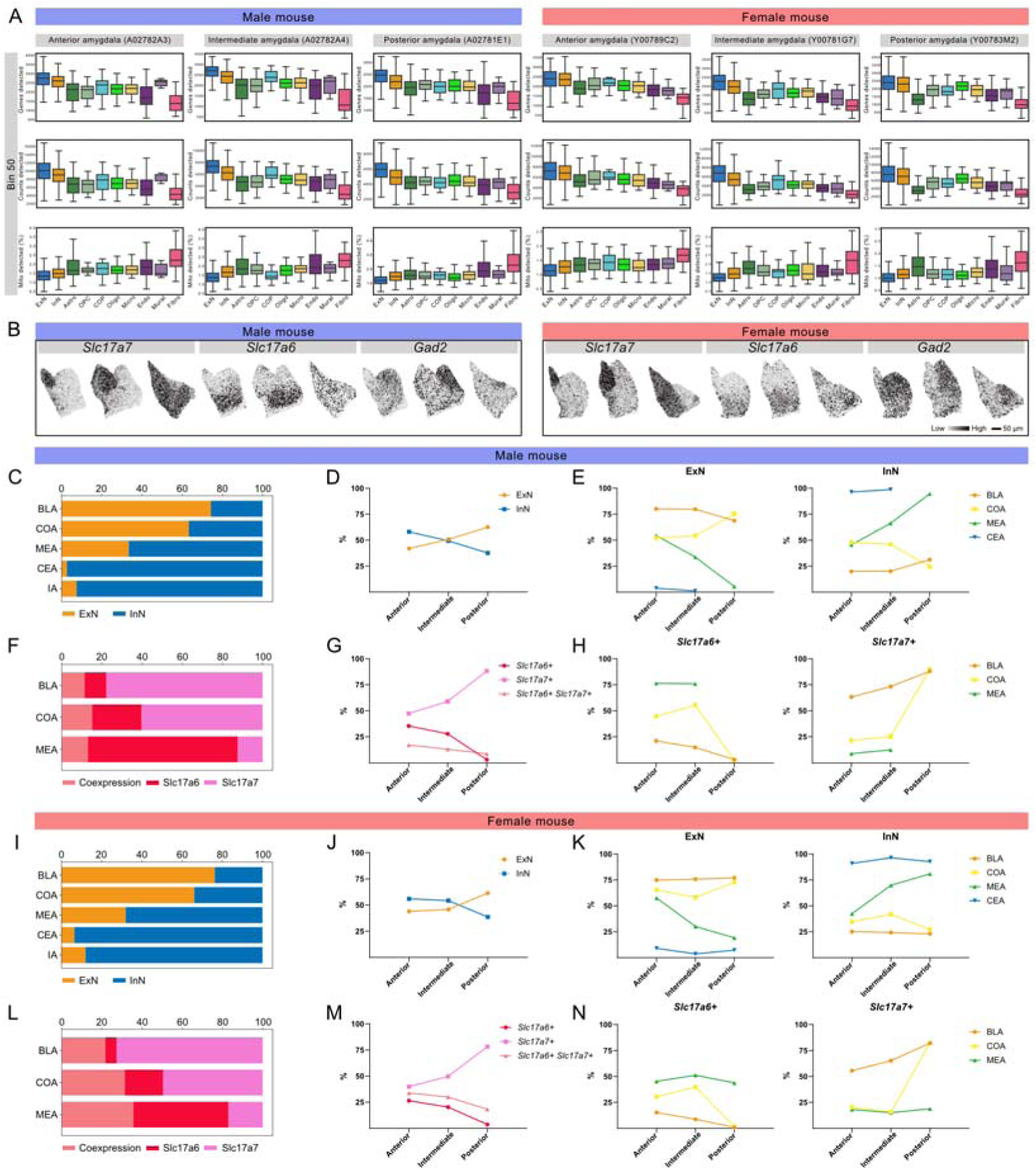
Quality control and spatial profiling of ExN, InN, and their marker genes in mouse amygdala, related to. Figure 2. (A) Quality control plots for Stereo-seq datasets of male and female mouse amygdala. Box plots showing total counts, detected genes, and mitochondrial genes per cell classes. Bin 50 data were shown; Chip IDs in parentheses. (B) Spatial expression patterns of *Slc17a7*, *Slc17a6*, and *Gad2* across anterior, intermediate, and posterior amygdala in male and female mice. Colors represent relative expression. (C and I) Proportions of ExN and InN across male (C) and female (I) amygdala subregions. (D and J) Proportions of ExN and InN along the anteroposterior axis of male (D) and female (J) amygdala. (E and K) Proportions of ExN and InN across anterior, intermediate, and posterior amygdala subregions in male (E) and female (K) mice. (F and L) Proportions of *Slc17a6*^+^, *Slc17a7*^+^, and double-positive neurons across male (F) and female (L) amygdala subregions. (G and M) Proportions of *Slc17a6*^+^, *Slc17a7*^+^, and double-positive neurons along the anteroposterior axis of male (G) and female (M) amygdala. (H and N) Proportions of *Slc17a6*^+^ and *Slc17a7*^+^ neurons across anterior, intermediate, and posterior amygdala subregions in male (M) and female (F) mice.

**Figure S3.**
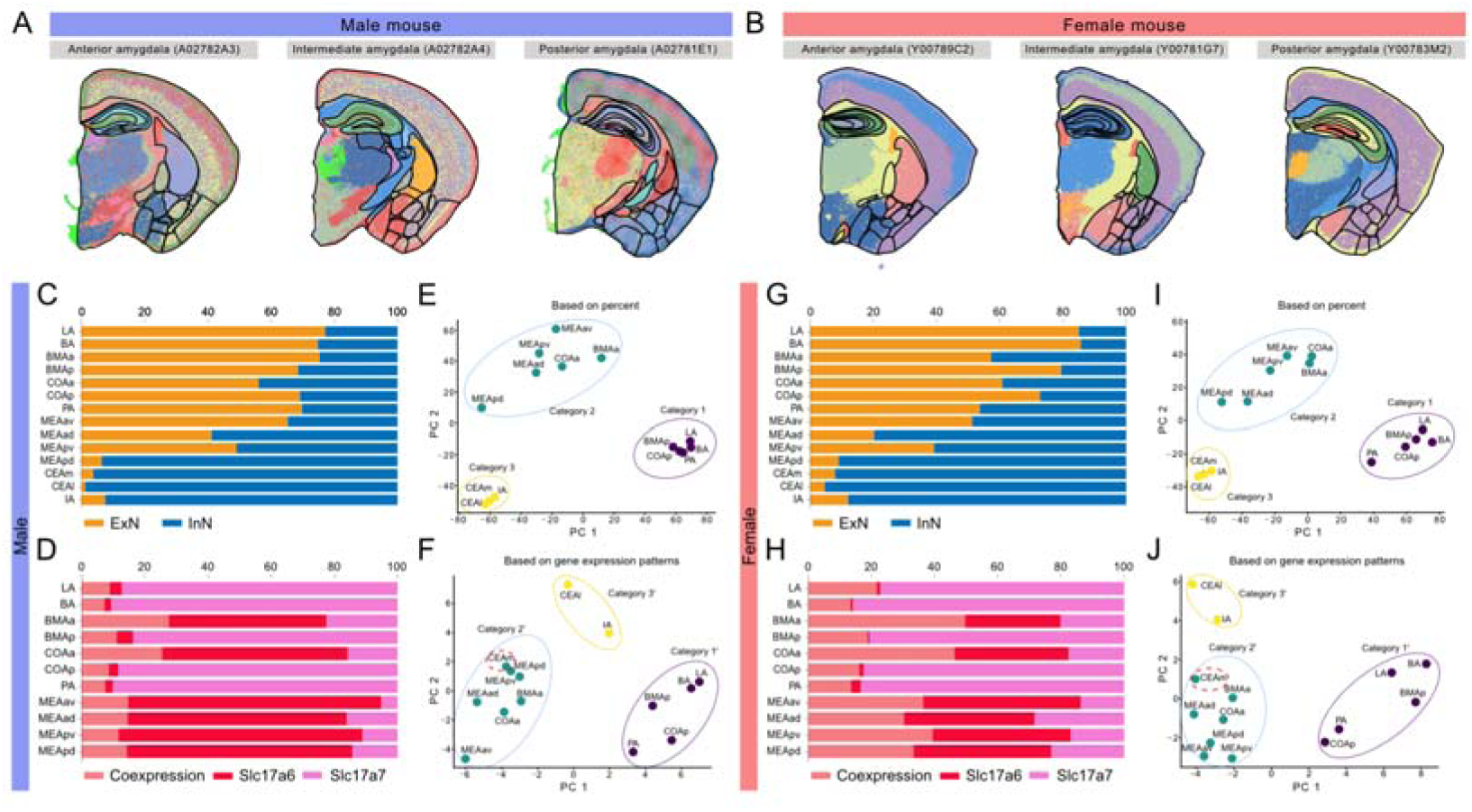
Subnuclear parcellation and novel subnuclear organization patterns in mouse amygdala, related to. Figure 2. (A and B) Stereo-seq data registered to Allen Mouse Brain Atlas for precise amygdala subnuclear parcellation in male (A) and female (B) mice. (C and G) Proportions of ExN and InN across male (C) and female (G) amygdala subnuclei. (D and H) Proportions of *Slc17a6*^+^, *Slc17a7*^+^, and double-positive neurons across male (D) and female (H) amygdala subnuclei. (E and I) PCA dimension reduction and clustering of male (E) and female (I) amygdala subnuclei by ExN, InN, *Slc17a6*^+^, *Slc17a7*^+^, and double-positive neuron proportions. Colors indicate subnuclei categories. (F and J) PCA dimension reduction and clustering of male (F) and female (J) amygdala subnuclei by feature gene expression. Colors indicate subnuclei categories’.

**Figure S4.**
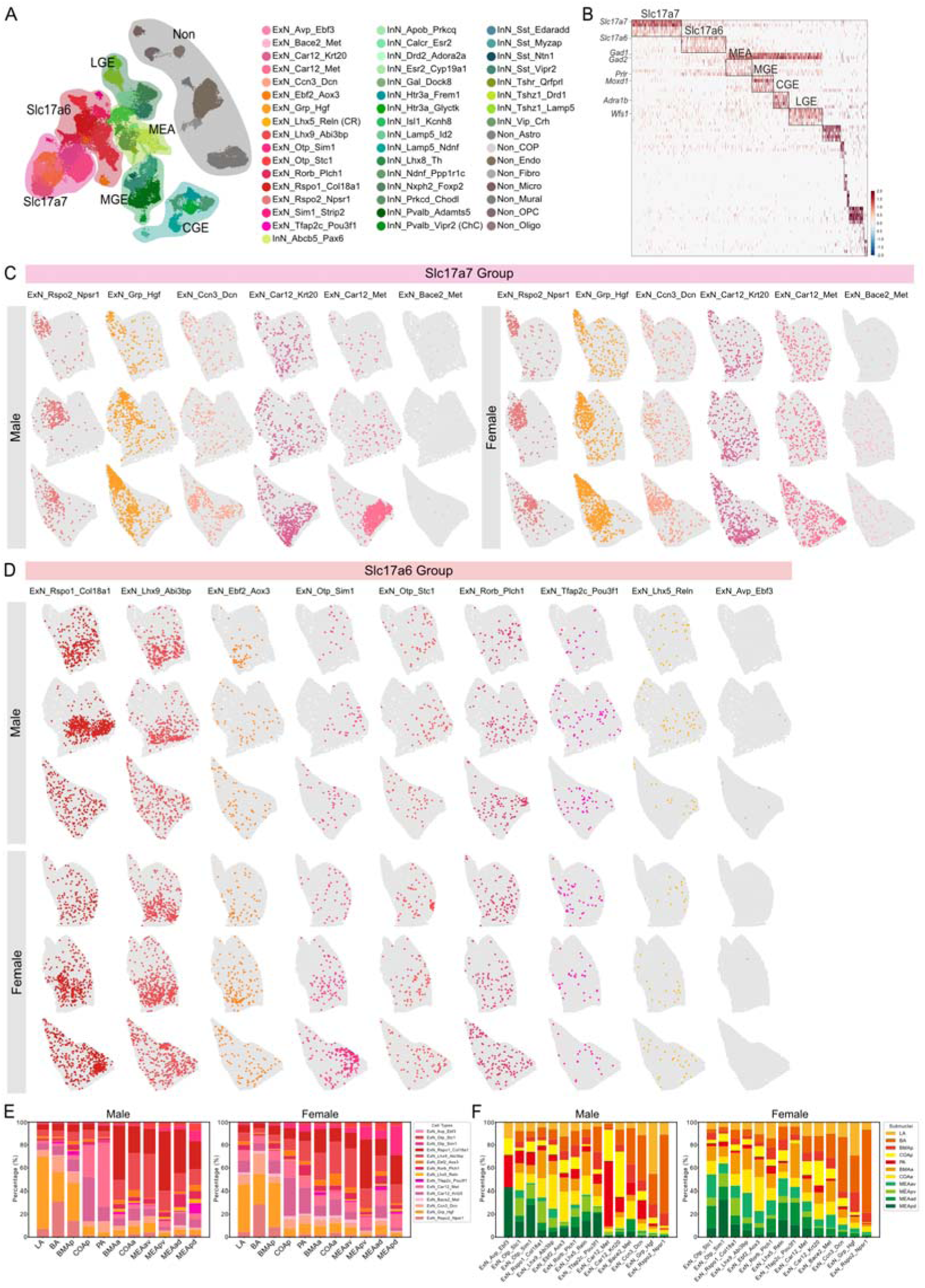
Stereo-seq reveals subnuclear distribution of excitatory neuron types at single-cell resolution, related to. Figure 3. (A) UMAP visualization of six neuronal groups in mouse amygdala, with distinct shading colors indicating each group. (B) Marker genes for six neuronal groups in mouse amygdala, with selected feature genes labeled. See Table S3. (C and D) Spatial distribution of Slc17a7 (C) and Slc17a6 (D) groups of neuron types in male and female mouse amygdala at single-cell resolution. (E) Proportions of excitatory neuron types across amygdala subnuclei in male and female mice. See Table S4. (F) Subnuclear distribution proportions of excitatory neuron types in male and female mice. See Table S4.

**Figure S5.**
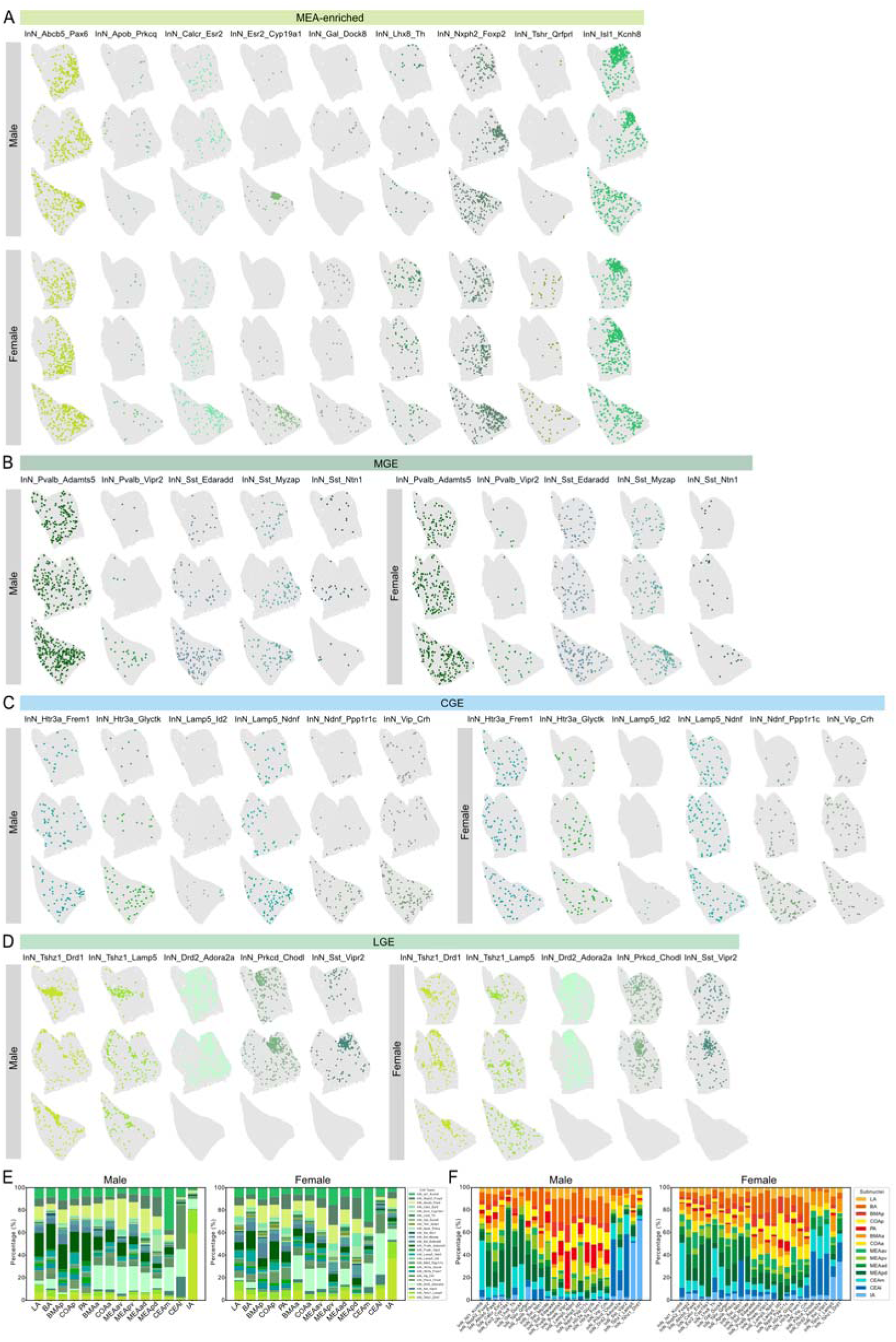
Stereo-seq reveals subnuclear distribution of inhibitory neuron types at single-cell resolution, related to. Figure 3. (A-D) Spatial distribution of MEA-enriched (A), MGE (B), CGE (C), and LGE (D) groups of neuron types in male and female mouse amygdala at single-cell resolution. (E) Proportions of inhibitory neuron types across amygdala subnuclei in male and female mice. See Table S4. (F) Subnuclear distribution proportions of inhibitory neuron types in male and female mice. See Table S4.

**Figure S6.**
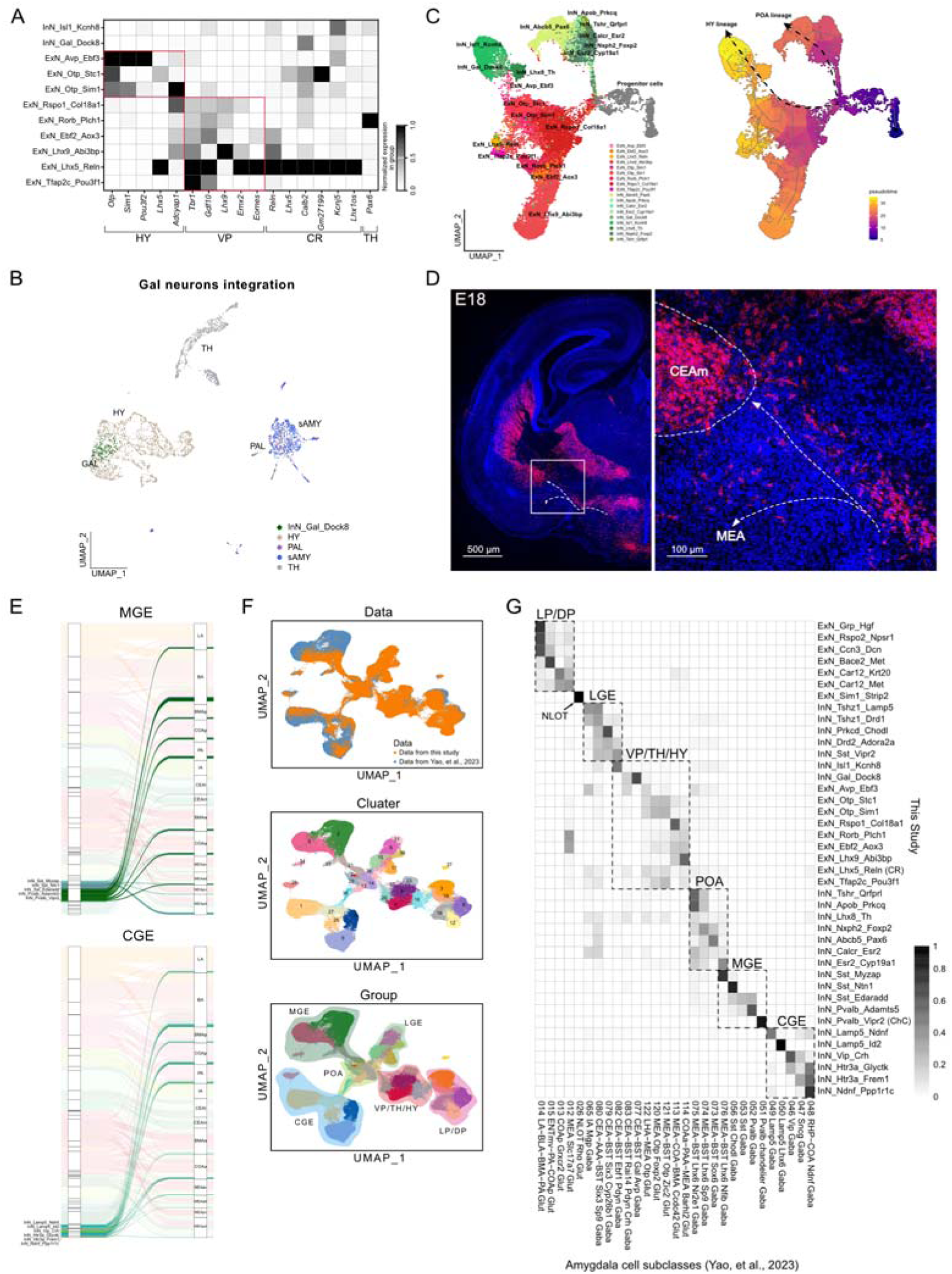
Developmental origins and subnuclear distribution of amygdalar neuron types in mice, related to. Figure 4. (A) Neuronal types in VP/TH/HY group expressing differential neurogenic niche marker genes. CR: Cajal-Retzius cells. Colors indicate normalized expression. (B) UMAP projection integrating InN_Gal_Dock8 with published *Gal*^+^ neurons across brain regions. (C) Neuronal types in VP/TH/HY and POA groups integrated with reported hypothalamic progenitor cells (left), with Monocle-inferred developmental trajectories (right). Arrows depict potential differentiation paths; pseudotime shown for individual cells (right). (D) RNAscope staining shows the distribution of *Isl1*^+^ neurons in the amygdala and adjacent regions at mouse embryonic day 18 (E18), as well as the migration pathways of hypothalamic-derived *Isl1*^+^ neurons (indicated by dashed lines with arrows). (E) MGE- and CGE-derived interneuron types are widely distributed across amygdalar subnuclei except CEAl, CEAm, and IA. (F) UMAP visualization integrating our mouse amygdala single-cell data with published subregion/subnucleus-annotated cell types. Nuclei colored by datasets (upper), and clusters (middle and bottom). Shaded areas indicate groups by developmental origin (bottom). (G) Proportion of nuclei overlapping between our amygdala and published amygdala clusters in integrated datasets. Dashed boxes highlight developmentally distinct groups.

